# Conformational landscape of the transcription factor ATF4 is dominated by disordered-mediated inter-domain coupling

**DOI:** 10.1101/2023.05.12.540518

**Authors:** Urval Patel, Steven Siang, Davit Potoyan, Julien Roche

## Abstract

Transient intramolecular interactions between transactivation domain and DNA binding domain of transcription factors are known to play important functional roles, including modulation of DNA binding affinity and specificity. Similar type of inter-domain interactions has recently been reported for the transcription factor ATF4/CREB-2, a key regulator of the Integral Stress Response. In the case of ATF4, transient coupling between the transactivation and basic-leucine zipper (bZip) domains regulates the degree of phosphorylation of the disordered transactivation domain achievable by the casein kinase CK2. Despite the crucial importance of these inter-domain interactions, their structural and molecular basis remain ill-determined. In the present study, we use a combination of experimental and computational techniques to determine the precise nature of the long-range contacts established between the transactivation and bZip domains of ATF4 prior to its association with protein partners and DNA. Solution NMR spectroscopy experiments reveal that the isolated bZip domain of ATF4 is predominantly disordered and display evidence of conformational dynamics over a wide range of timescales. These experimental findings are supported by multi-microsecond timescale all-atom molecular simulations that unveil the molecular basis of the long-range interactions between the transactivation and bZip domains of ATF4. We found that inter-domain coupling is primarily driven by disorder-mediated interactions between a leucine-rich region of the transactivation domain and the leucine-zipper region of the bZip domain. This study uncovers the role played by structural disorder in facilitating the formation of long-range intramolecular interactions that shape the conformational ensemble of ATF4 in a critical manner.

## INTRODUCTION

Activating transcription factor 4 (ATF4) is a basic leucine zipper (bZip) transcription factor that plays a central role in the Integral Stress Response (ISR) by communicating pro-survival and pro-apoptotic signals in response to extracellular stresses.^1-3^ When cells suffer from various stresses, such as amino acid deprivation, oxidative stress, or hypoxia, ATF4 is transcriptionally and translationally upregulated in response to eIF2α phosphorylation.^4-6^ ATF4 can form heterodimers with a range of other bZip transcription factors, each heterodimer controlling a specific array of regulatory mechanisms and generating a unique set of downstream effects (**Fig. 1A**).^7^ Regulatory mechanisms such as post-translational modifications enable ATF4 to direct transcriptional expression of a very large number of stress-related genes, often as an activator, sometimes as a repressor.^2^ Such wide adaptive control is enabled by the predominantly disordered nature of ATF4, which is essential for establishing optimal adaptation that is tailored toward different stress signals. Therefore, it is important to precisely determine the role played by structural disorder in adaptive control to understand the implementation of stress-adaptive transcriptomes in healthy and disease-related conditions.

**Figure 1.**
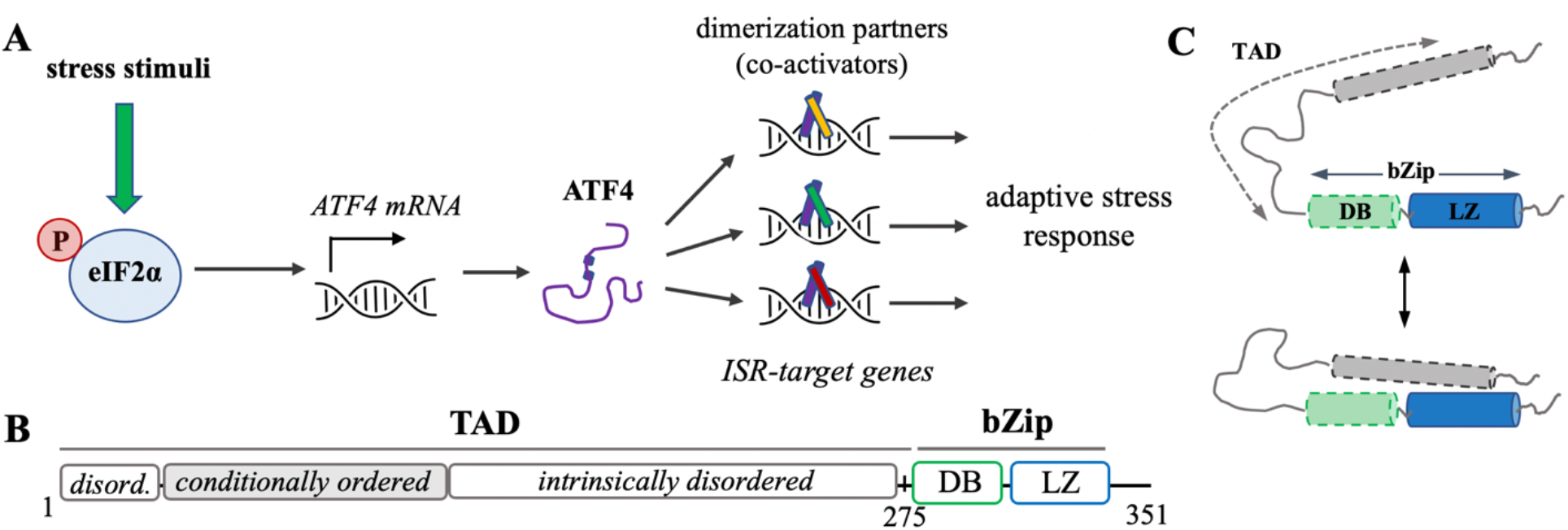
**(A)** Schematic representation of ATF4 activation in response to stress stimuli. Various stress stimuli can promote the phosphorylation of eIF2α, the central mediator of the ISR. Activation of eIF2α leads to the transcriptional and translational upregulation of *ATF4*. ATF4 can associate with a wide range of bZip co-activators to control the activation (or repression) of ISR-target genes. (**B**) Representation of ATF4 sequence showing the N-terminal transactivation domain (**TAD**, residue 1-275) and the C-terminal basic-leucine zipper domain **bZip** domain, which is composed of a DNA binding region (**DB**, residues 280-301) and a leucine-zipper region (**LZ**, residues 306-344). Here the LZ region is shown as intrinsically ordered, the DB and first half of the TAD as conditionally ordered, while the second half of the TAD is known to be intrinsically disordered. (**C**) Illustration of the transient interactions taking place between the TAD (gray) and bZip domain composed of the DNA binding region (green) and leucine zipper region (blue).

ATF4 contains a C-terminal bZip domain composed of a DNA binding (DB) region rich in basic residues (residues 280-301) and a leucine zipper (LZ) region (residues 306-334) essential for dimerization (**Fig. 1B**).^3^ The DNA binding region of bZip transcription factors is typically found to be predominantly disordered and to experience fast conformational dynamics in the absence of DNA, while the zipper region is usually more stable and more structured.^8,9^ The rest of ATF4 consists of a transactivation domain (TAD, residues 1-275) that contains multiple post-translational modification sites and mediates interactions with numerous partner proteins.^10^ The transactivation domain is known to regulate ATF4 stability and localization and its capacity to modulate gene transcription.^11^ Our group has recently conducted an in-depth investigation of the structural properties of the transactivation domain of ATF4 in a monomeric state using a combination of solution NMR spectroscopy, SEC-MALS, and SAXS.^12^ Backbone ^13^C chemical shifts and temperature coefficients point to the TAD being largely disordered in solution, with the exception of small helical motifs at the N-terminal end of the domain. Remarkably, when the NMR spectrum of the isolated TAD was compared with that of the full-length ATF4, we observed no additional amide crosspeak that could be assigned to the bZip domain.^12^ This suggests that residues of the bZip domain experience conformational exchange on an intermediate timescale that leads to excessive line-width broadening. Furthermore, titration of ^15^N-labeled isolated TAD with unlabeled isolated bZip domain shows significant attenuation of crosspeaks within the N-terminal region of the TAD. All together, these results indicate that in the monomeric state the TAD and bZip domains of ATF4 are not structurally independent but form transient long-range interactions in solution.^12^ These interactions involve the first N-terminal half of the TAD that is predicted to be conditionally ordered and the leucine zipper region of bZip (**Fig. 1C**). The presence of the bZip domain was also found to be required in order to achieve phosphorylation of TAD by the kinase CK2. Long-range interactions between the TAD and bZip domains of ATF4 appear therefore to modulate the accessibility of the TAD to a protein kinase. Given the prominent role played by misregulation of ATF4 in various diseases such as cancer^13^, muscle cachexia^14^, liver disease^15^, and cardiovascular disease^16^, these results open new perspectives for targeting on-pathway monomeric states of ATF4 prior to its association with protein partners or DNA. The goal of such strategy is to disrupt specific sets of intramolecular interactions that play a key role in shaping the conformational ensemble of ATF4.

In the present study, we seek to characterize the molecular mechanisms underlying the formation of long-range interactions between the TAD and bZip domains of ATF4 using a combination of biophysical techniques and all-atom molecular dynamics simulations. Characterization of the isolated bZip domain of ATF4 revealed a high degree of conformational disorder in the absence of DNA, which suggests that the full-length ATF4 is predominantly disordered prior to its association with other bZip partners and/or DNA. Simulation of the full-length protein revealed that ATF4 preferentially samples an ensemble of structurally disordered but compact conformations dominated by long-range interactions between the TAD and bZip domains. This study provides new insights into the disorder-mediated mechanisms responsible for establishing long-range interactions between the TAD and bZip domains, which is critical for understanding the mechanisms of action of ATF4.

## RESULTS

### 1. Biophysical characterization of isolated bZip domain

Since excessive linewidth broadening due to conformational exchange prevents the direct observation of amide resonances from bZip residues in the full-length ATF4 spectra, we expressed and purified the isolated bZip domain for further study by solution NMR. CD spectra recorded in the absence of DNA show that the isolated bZip is predominantly disordered in solution, in good agreement with previous reports indicating that the basic region of bZip domains is typically disordered in the absence of DNA (**Fig. 2A**).^8,9,17^ We observed a significant change when bZip is mixed with the double-stranded CRE sequence (18 base pairs) indicating that the isolated bZip domain is able to bind to DNA and acquires a more helical conformation upon binding. We further characterized the isolated bZip construct using solution NMR spectroscopy. 2D ^1^H-^15^N NMR spectra collected for the isolated bZip domain show that more than 50% of the expected crosspeaks were observed both in the absence as well as in the presence of DNA (**Fig. 2B**). It indicates that residues of both the free and DNA-bound bZip domain experience excessive linewidth broadening due to conformational exchange on an intermediate timescale. Partial backbone chemical shift assignment revealed that the observable resonances correspond to amino acids located mainly in the DNA binding region and in the linker connecting the DB and LZ regions (in gray in **Fig. 2B** upper panel). The backbone ^13^C chemical shifts of those residues show minimal deviations from random coil in the absence of DNA, indicating that this portion of bZip is predominantly disordered in the DNA-free form (**Fig. S1**). Triple resonance experiments were also recorded in the presence of DNA, unfortunately chemical shifts assignment was hampered by sample stability issues and excessive linewidth broadening.

**Figure 2:**
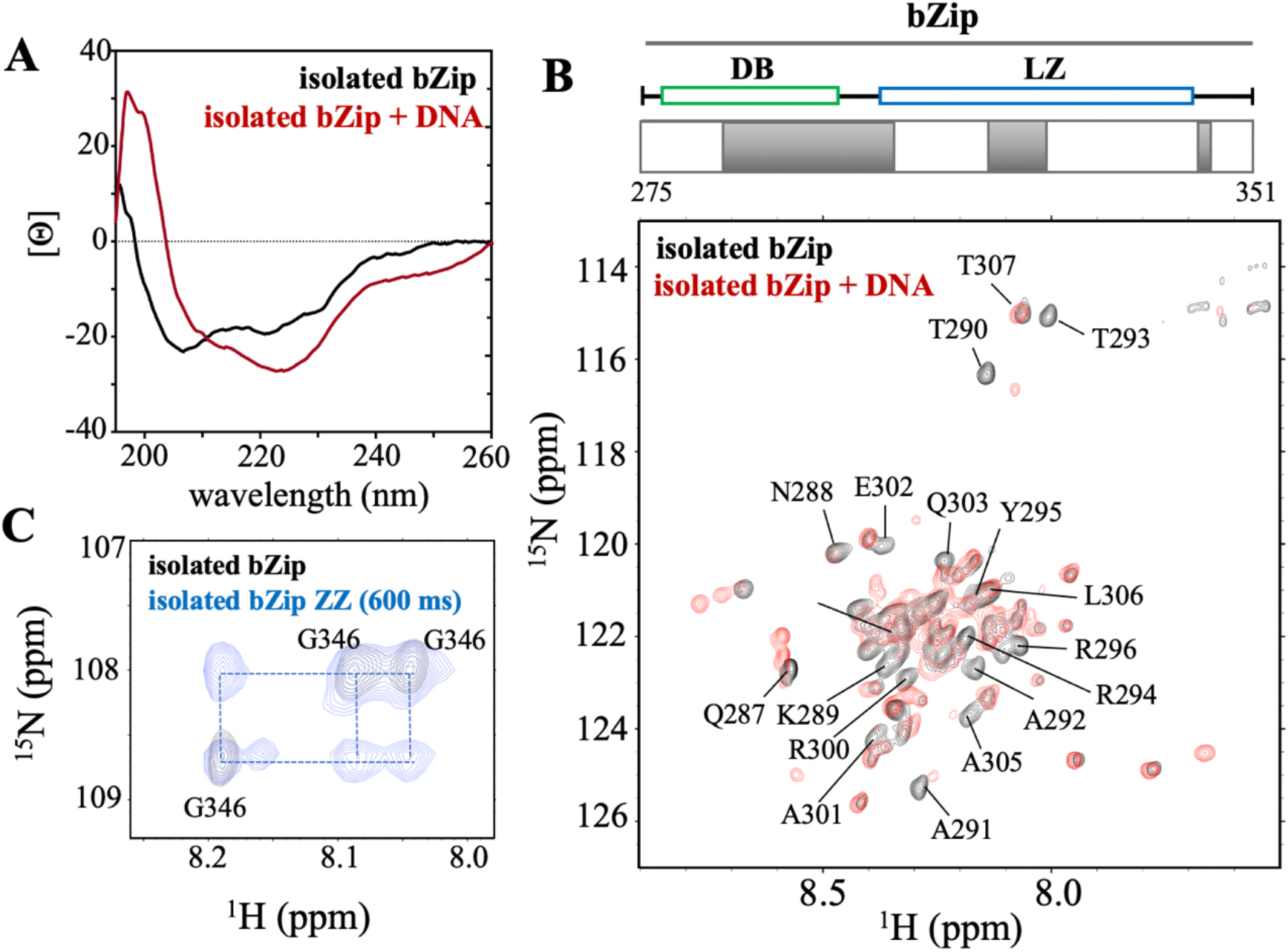
**(A)** CD spectra collected at 293 K for the isolated bZip domain in the absence (black) and in the presence (red) of double-stranded DNA corresponding to the CRE consensus sequence (18 base pairs) at equimolar ratio. (**B**) (*upper panel*) Schematic representation of the bZip domain with the DNA binding regions (**DB**) shown in green and the leucine zipper region (**LZ**) in blue. Portions of the bZip domain for which we were able to assign the backbone chemical shifts are shown in gray. (*lower panel*) 2D ^1^H-^15^N spectra of the isolated bZip domain in the absence (black) and in the presence (red) of double-stranded CRE motif at equimolar ratio. (**C**) Small region of 2D ZZ-exchange experiments collected for the isolated bZip domain in the absence of DNA. Three major crosspeaks of similar intensities were assigned to the C-terminal glycine G346 (black). Buildup of minor crosspeaks was observed for ZZ-exchange experiments with a mixing time of 600 ms (blue), indicating that G346 is experiencing chemical exchange between at least three distinct conformational states on a slow timescale relative to NMR timescale.

Interestingly, further investigation into the conformational dynamics of the bZip domain in the absence of DNA revealed that the isolated bZip domain experiences a broad range of dynamical regimes. Indeed, we noticed for instance three amide resonances of similar intensity assigned to the C-terminal glycine G346 (**Fig. 2C**, black spectrum). Evidence of a slow exchange process can be observed in ZZ-exchange experiments that show build-up of minor peaks for the three sets of amide resonances (**Fig. 2C**, blue spectrum). It indicates that G346, and potentially a significant part of the C-terminal region of bZip, is in exchange between at least three different conformational states. Overall, these results suggest that the bZip domain is experiencing conformational exchange over a wide range of timescales, from intermediate (μs) to slow (μs-ms) relative to NMR timescale.

### 2. All-atom MD simulations of the isolated bZip domain

A series of independent all-atom MD simulations were then run to further characterize the structure and dynamics of the partially disordered bZip domain. The simulations were conducted with AMBER99SB-disp, a force field that combines a newer generation of water model with a set of parameters specifically designed to capture the behavior of partially disordered proteins in solution.^18^ Starting from a fully helical conformation of the bZip domain extracted from the structure of the ATF4-c/EBP heterodimer (PDB 1CI6)^17^, 4 independent all-atom MD simulations of the isolated bZip domain of ATF4 were run for a combined total production run of 8 μs. Analysis of secondary structure content with DSSP as well as principal component analysis (PCA) in the ϕ, φ dihedral space reveal that the initial helical conformation converges toward a predominantly disordered ensemble following different unfolding pathways. The DNA binding region of the bZip domain appears to be less stable than the leucine zipper region in trajectories 1 and 4, while the opposite was observed in trajectories 2 and 3 (**Fig. S2**), suggesting that the DNA binding region and leucine zipper are loosely coupled and experience independent unfolding pathways. Regardless of the unfolding pathway adopted, all four trajectories reach a predominantly disordered state after ∼1.2 μs. This initial set of simulations confirmed that in isolation the monomeric bZip domain of ATF4 is predominantly disordered in solution, in good agreement with the CD and NMR data described above. In order to further sample the disordered conformational ensemble adopted by the bZip domain, two additional all-atom simulations starting from a representative disordered conformation were run for a total of 10 μs. PCA conducted in the ϕ, φ dihedral space reveals a complex conformational landscape with multiple basins of the population (**Fig. 3A**). Representative conformations extracted from the three most populated basins through cluster analysis point to a predominantly disordered structural ensemble with the presence of short helical motifs predominantly located in LZ region (insert **Fig. 3A**). DSSP analysis conducted on the combined 10 μs trajectories reveals that the LZ region displays indeed a significantly higher propensity for helical conformations than the DB region (**Fig. 3B**). Interestingly, the short linker between the two regions (residue 302-305) shows a relatively high helical propensity (∼60% for E304), suggesting that this linker may potentially help with the folding of the DB region in appropriate conditions. Altogether, the combination of experimental evidence (CD and NMR) and computational data paints a picture of the isolated, monomeric bZip domain of ATF4 being largely disordered in solution, with a partially helical leucine zipper region and a predominantly disordered DNA binding region.

**Figure 3:**
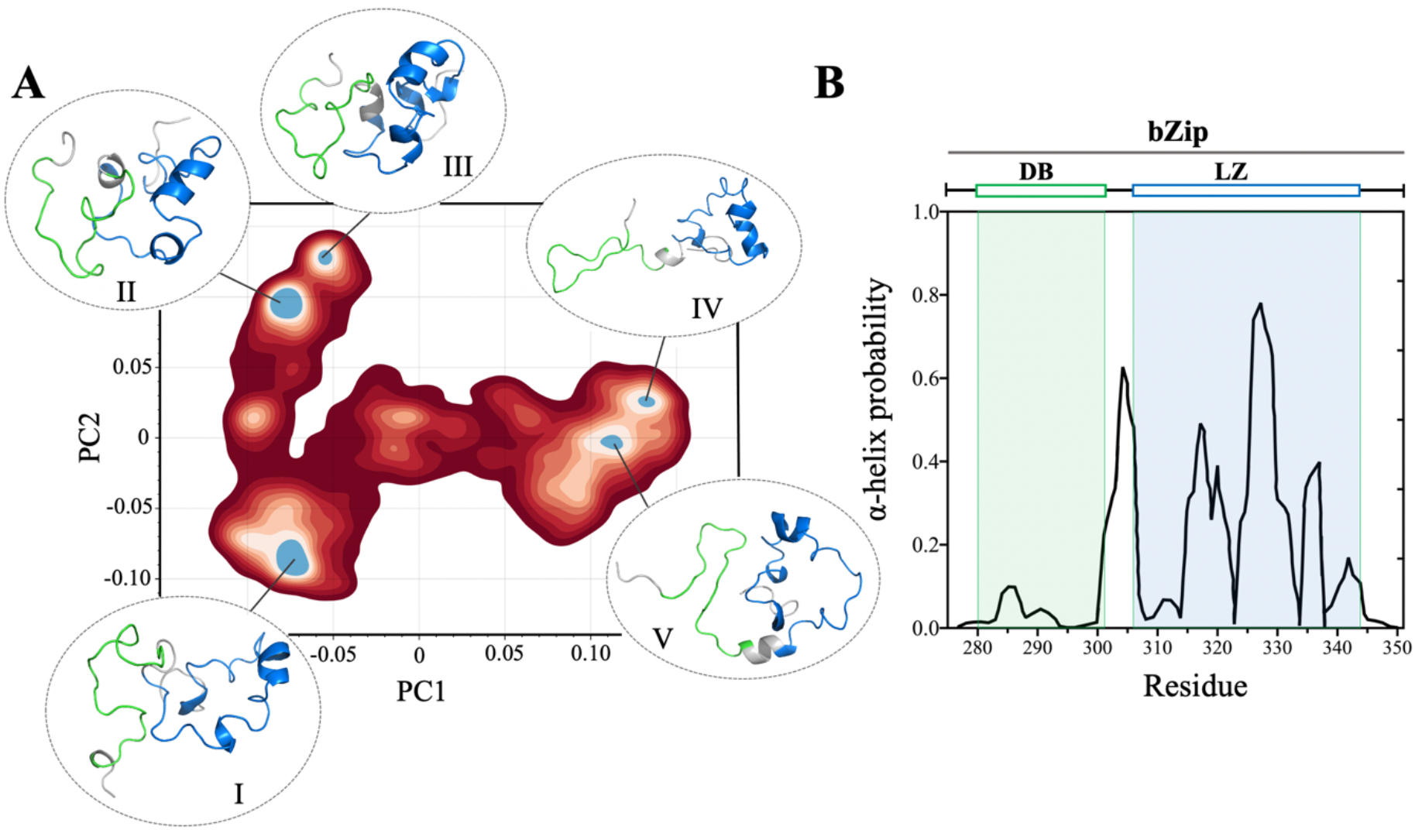
**(A)** Probability distribution of isolated bZip domain conformations (residue 276-351) generated through 10 μs all-atom simulations and projected onto the first two principal components of the ϕ, φ dihedral space (blue: high-probability, dark red: low probability). Inserts show representative conformations extracted from the five most populated basins (I-V). The DNA binding region (DB) is colored in green and the leucine zipper region (LZ) is colored in blue. (**B**) Residue-specific alpha-helical propensity calculated from DSSP analysis of the combined 10 μs all-atom simulations. The DB and LZ regions are highlighted in green and blue respectively.

### 3. All-atom MD simulations of the full-length ATF4

Next, we conducted long timescale all-atom MD simulations of the full-length ATF4 in order to determine the nature and molecular mechanisms involved in the long-range interactions between the TAD and bZip domains that have been recently reported. Three independent replicas were set up with AMBER99SB-disp using a partially disordered conformation of the monomeric ATF4 generated by RaptorX^19^ as starting conformation. Using PCA in the Cα-Cα contact space, we observed convergence of the three independent replicas within 900 ns (**Fig. S3**). Each trajectory was further extended to 4 μs to ensure sufficient sampling for a combined total production run of 9.3 μs. Conformation frequency distribution projected onto the first two PC in the Cα-Cα contact reveals a landscape with relatively low conformational heterogeneity. Indeed, a single major basin represents nearly 80% of all conformations (**Fig. 4A**). Cluster analysis conducted on this basin shows that the predominantly populated conformations are compact and partially disordered (insert, **Fig. 4A**). Interestingly, the bZip domain appears more structured in the full-length ATF4 than in isolation. This is especially true for the LZ region of the bZip domain, which is almost fully helical in the context of the full-length ATF4 but at most ∼60% helical in the isolated bZip domain (**Fig. 3B**). The stabilization of the bZip domain and the overall compactness of the full-length ATF4 is explained by an extensive network of long-range contacts between the TAD and bZip domains as displayed in the frequency Cα-Cα contact map (**Fig. 4B**). Notably, the N-term region of the TAD (residue 102-125) is found to establish a large number of contacts with the LZ region (residue 300-347), in excellent agreement with the recently published NMR data. The Cα-Cα contact map reveals indeed that most frequent long-range contacts between the TAD and bZip domains are established between a leucine-rich region of the TAD encompassing L105, L112, L113, L116, and L122, and the leucine-zipper region of the bZip domain via L320, L327, and L334 (**Fig. 5**). These hydrophobic interactions are frequently complemented by two ionic bonds formed between D110-R323 and D90-R342 (**Fig. 5**).

**Figure 4:**
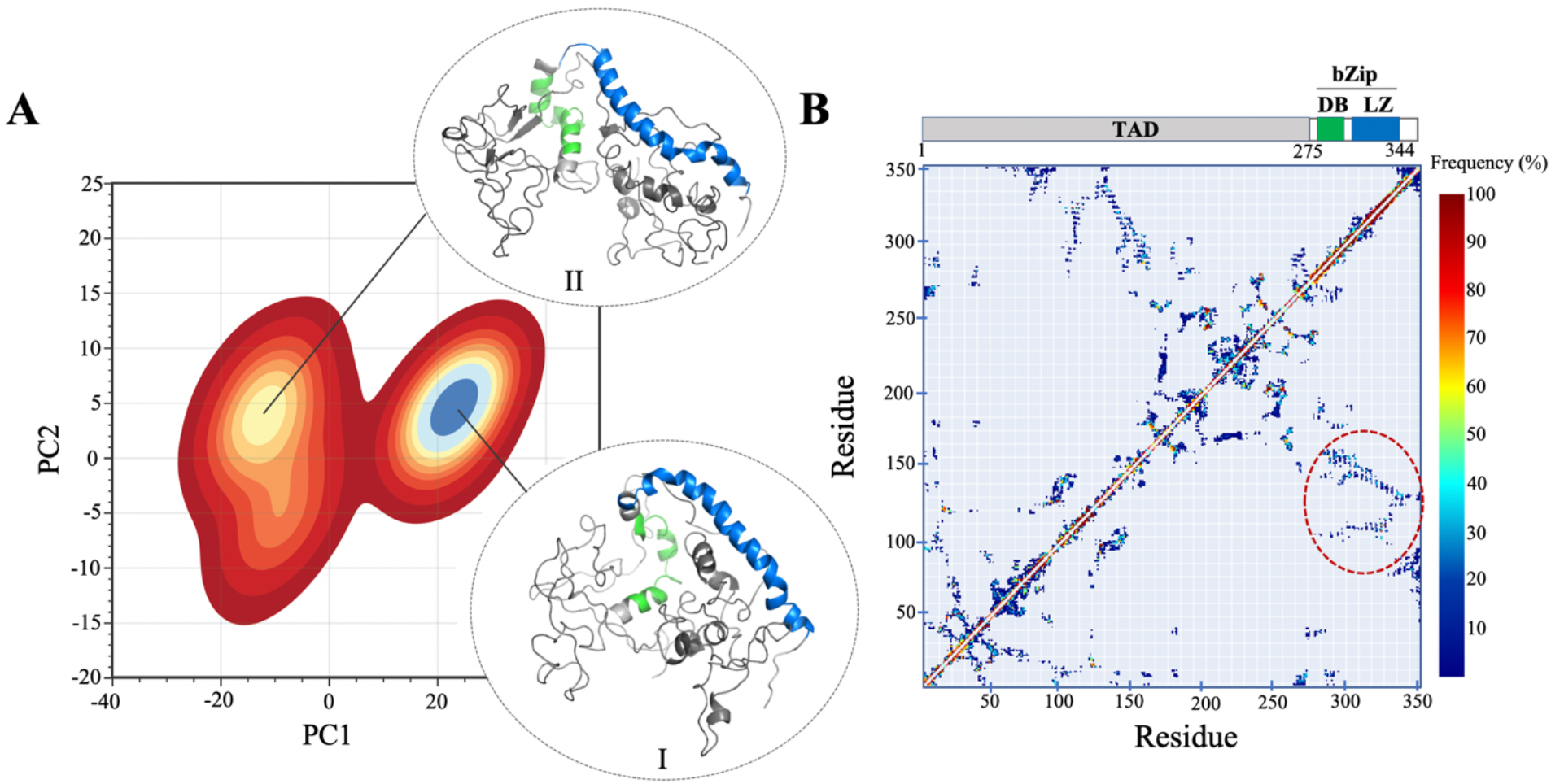
**(A)** Probability distribution of full-length ATF4 conformations generated through 4.8 μs all-atom simulations and projected onto the first two principal components of the Cα-Cα contact space (blue: high-probability, dark red: low probability). Inserts show representative conformation extracted from the most populated basin. The DNA binding region (DB) is colored in green and the leucine zipper region (LZ) is colored in blue. The TAD domain is shown in dark gray. (**B**) Cα-Cα frequency contact map calculated over 4.8 μs all-atom simulation of full-length ATF4. The contact reveals an extensive network of long-range interactions between the N-terminal region of the TAD (residues 102-155) and the LZ region of the bZip domain (residues 300-347) (highlighted with a dashed red circle).

**Figure 5:**
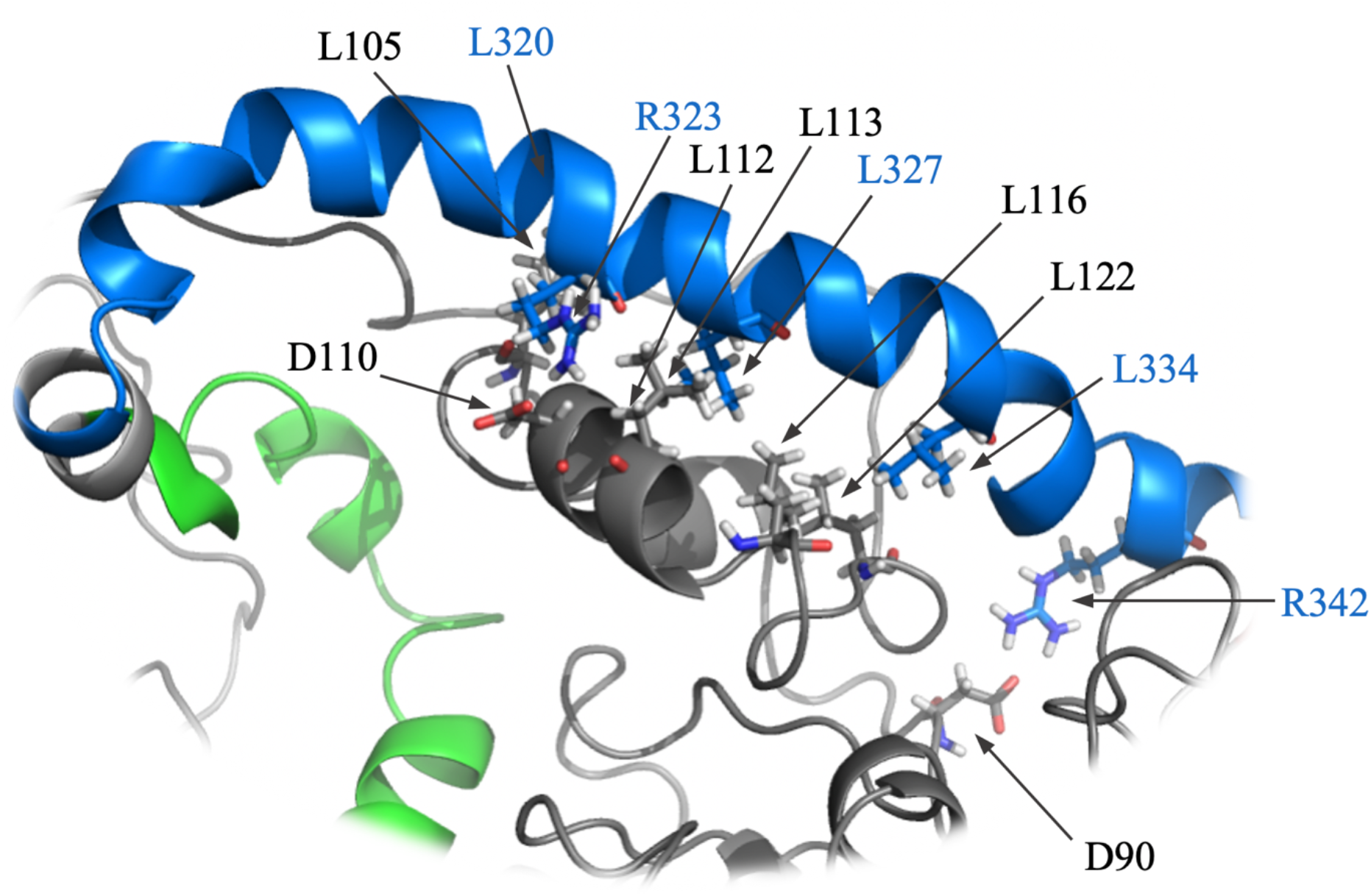
Close-up of a representative conformation of the full-length ATF4 isolated from the main population basin determined by PCA (**Fig. 4A**), highlighting the most frequent contacts established between the TAD and bZip domains. These contacts involve a leucine-rich region located at the N-terminus of the TAD (L105, L112, L113, L116, and L122) and the leucine-zipper region of the bZip domain (L320, L327, L334). Hydrophobic contacts between leucines are frequently found to be further stabilized by ionic bonds such as D110-R323 and D90-R342.

## DISCUSSION

Misregulation of transcription factors is associated with a broad range of human diseases, including cancers, autoimmune and neurological disorders, diabetes, and cardiovascular disease.^20-24^ Yet despite their immense potential, transcription factors are often considered an undruggable protein class.^25,26^ Historically, this has been attributed to the lack of specificity of transcription factor interaction networks (i.e. a large number of different transcription factors can share identical protein partners) and the disordered nature of transcription factors that makes traditional drug discovery approaches ill-suited. Therapeutic strategies explored over the past years include (i) disrupting interaction with their binding partners^27^, (ii) targeting well-structured regulatory domains of transcription factors^28^, and (iii) blocking formation of transcription factor-DNA complexes.^29^ The aim of the present study is to pave the way toward a different strategy that consists in targeting on-pathway monomeric states of transcription factors prior to their association with protein partners and DNA. As a canonical member of the bZip family of transcription factors, ATF4 constitutes an excellent model for studying the nature of disorder-mediated intramolecular interactions that may be critical for the formation of functional dimeric complexes.

Long-range couplings between the transactivation and DNA binding domains have been reported for multiple transcription factors, including non-bZip transcription factors such as p53^30^ and FOXO4.^31^ In most cases, such coupling appears to modulate the binding affinity of the DNA binding domains to specific promotor regions. The binding affinity to DNA can be further tuned by post-translational modifications of the transactivation domain (typically phosphorylation).^32-35^ A similar mechanism of inter-domain coupling has been recently reported for the transcription factor ATF4. In the case of ATF4, we found that long-range coupling between the TAD and bZip domains plays a critical role in modulating the accessibility of the TAD to the protein kinase CK2.^12^ In the present study, we aim to determine the precise molecular mechanisms underlying the formation of the transient interactions between the TAD and bZip domains of ATF4.

We first examined the conformational properties of the isolated bZip domain in solution. 2D ^1^H-^15^N NMR spectra recorded on the truncated bZip domain show evidence of extensive conformational exchange across multiple timescales, from intermediate to slow relative to NMR timescale, especially affecting residues within the leucine zipper region (**Fig. 2 B,C**). Partial backbone chemical shift assignment revealed that the DNA binding region is predominantly disordered in such conditions (**Fig. S1**). These findings are reminiscent of other bZip domains such as GCN4, the budding yeast homologue of ATF4. Palmer and coworkers have indeed demonstrated that the DNA binding region of GCN4 is predominantly disordered and experiences fast conformational dynamics in the absence of DNA, while the zipper region is more stable and experiences conformational exchange on a slower timescale.^8,9^ We then conducted a series of long-timescale all-atom molecular dynamics simulations to gain further insights into the structural features of the isolated bZip domain. Principal component analysis conducted on the ϕ, φ dihedral space reveals a complex, multiple-basin conformational landscape (**Fig. 3A**). In good agreement with the NMR data, these simulations show that the DNA binding region is predominantly disordered while the leucine zipper region is partially helical (**Fig. 3B**). This combination of biophysical and computational characterization suggests that in isolation the bZip domain is, similarly to other bZip domains, largely disordered in solution. Altogether, the present study and the recent NMR analysis conducted on TAD outline the high propensity of both functional domains of ATF4 for structural disorder.

Next, we examined the structural features of the TAD and bZip domains in the context of the full-length ATF4 based on long-timescale all-atom simulations. In excellent agreement with the recently reported NMR data^12^, the MD simulations reveal an extensive network of long-range interactions between the TAD and bZip domains (**Fig. 4**). The inter-domain contacts involve the N-terminal region of the TAD (residues 105-151) and the leucine-zipper region of the bZip domain (**Fig. 4B**). These interactions significantly increase the helical content of the involved regions compared to their isolated counterparts, suggesting an induced-folding (or possibly conformational selection) mechanism of interaction. Notably, we found that the most frequent contacts involve a leucine-rich segment of the N-terminal region of the TAD (**Fig. 5**), which has been previously identified through NMR titration experiments conducted with ^15^N-labeled TAD and unlabeled bZip.^12^ Bioinformatic analysis have shown that the sequence composition of N-terminal region of the TAD differs from canonical intrinsically disordered sequences. The N-terminal region contains indeed significantly less hydrophilic and disorder promoting amino acids (Pro, Ser, and Thr) compared to the C-terminal region of the TAD.^12^ This further supports the potential formation of transient helical motifs within the N-terminal of the TAD upon interaction with the leucine-zipper region of the bZip domain. Finally, it is noteworthy that two amino acids found to be involved in frequent long-range contacts with the bZip domains (D110 and L112, **Fig. 5**) are listed in the Sanger database (https://cancer.sanger.ac.uk) as sites for nonsynonymous mutations isolated in cancer patients, which suggests a potential pathological significance of the long-range couplings between the TAD and bZip domains of ATF4.

Altogether, these findings highlight the nature of the long-range inter-domain interactions that shape the conformational ensemble of ATF4 prior to its association with protein partners or DNA. We found that these disorder-mediated contacts are primarily taking place between a leucine-rich region at the N-terminus of the TAD and the leucine-zipper region of the bZip domain. Ongoing work in our group investigates whether similar type of inter-domain coupling are shared by other members of the ATF family of transcription factors. Disrupting coupling between the TAD and bZip domains using therapeutic peptides or peptidomimetics can open new avenues for inhibiting the activity of this key family of transcription factors.^36^

## EXPERIMENTAL PROCEDURES

### Sample preparation

cDNA fragment covering the bZip domain of ATF4 (275-351) was amplified by PCR and inserted to pGEX-4T1 using EcoRI and XhoI restriction sites. TEV protease cut site was introduced between N-terminal glutathione S-transferase (GST) tag and bZip domain. A single point mutation (C310A) was introduced by PCR mutagenesis to improve stability at high sample concentration. The construct was transformed into E. coli protein expression strain BL21 (DE3) and protein expression was induced at an OD_600_ of 0.8 with 0.4 mM IPTG at 16°C overnight. Cells were harvested by centrifuging at 6,000 rpm and resuspended in lysing buffer (50mM CHES 9.0, 100mM NaCl, 5mM DTT, 1mM EDTA) in the presence of PMSF, Leupeptin, DNase I, and Halt Protease Inhibitor Cocktails (Thermo Scientific). Cells were then lysed by sonicating at 4 °C. Purification was initiated by passing the supernatant through Glutathione-Sepharose beads (UBPBio). Unbound proteins and contaminated DNA/RNA were washed with 50 mM CHES pH 9.0, 500 mM NaCl, 5 mM DTT, and 1 mM EDTA buffer solution. Cleaved GST-tag and bZip construct were separated by gel filtration chromatography using Hi-Load 16/600 Superdex 200 pg (GE Healthcare) on AKTA pure in a 20 mM MES pH 6.0, 50 mM NaCl, 1 mM EDTA buffer solution. All purification steps were carried on at 4 °C. Complementary oligonucleotides containing CRE motif (5′; - GCAGATGACGTCATCTGC - 3′) purchased from ISU DNA facility were resuspended in water to 1mM. DNA concentrations were determined by measuring the absorbance at 260 nm on NanoDrop spectrometer (Thermo Scientific). Equal volumes of complementary oligonucleotides were mixed and pelleted using a vacuum concentrator. dsDNA was synthesized by adding CD buffer, heating at 95°C for 10 minutes, and cooling it to 25°C slowly using a thermal cycler.

### Circular Dichroism

Circular dichroism (CD) spectra of isolated bZip (with/without DNA) at 20 μM were recorded using a MOS-500 (Biologic) spectropolarimeter controlled with Bio-Kine 32 at room temperature in a 1 mm UV quartz cuvette (FireflySci). The mean spectra were derived from three accumulations in the range of 190-260 nm. All data acquisitions were set for 0.2 sec with 2 nm slit width. Spectra were background-corrected with CD buffer (20 mM MES 6.5, 25 mM NaCl, 1 mM EDTA).

### Solution NMR Spectroscopy

NMR spectra were collected at ISU Biomolecular NMR Facility on a Bruker 700 MHz and 800 MHz spectrometers, equipped with z-shielded gradient triple resonance 5 mm TCI cryoprobe. NMR samples (either ^1^H/^15^N or ^2^H/^15^N/^13^C labeled) were prepared at protein concentrations of 200 μM in 50 mM potassium phosphate pH 6.0, 25 mM NaCl, 10 mM MgCl_2_, 1 mM EDTA, and 90% H_2_O/10% D_2_O (v/v). 2D ^1^H-^15^N TROSY-HSQC experiments were recorded with a time domain matrix consisting of 100* (t_1_, ^15^N) × 1024* (t_2_, ^1^H) complex points with acquisition time of 104 ms (t_1_) and 121 ms (t_2_) using 1.5 s interscan delay. Spectral widths for ^1^H and ^15^N dimensions were set to 15.9 and 28.2 ppm respectively, with carriers set at 4.821 ppm (^1^H) and 119.138 ppm (^15^N). 2D ^1^H-^15^N experiments conducted with dsDNA CRE motif were carried on at 800 MHz using an equimolar ^15^N-labeled bZip: DNA ratio. For these experiments a truncated version of the isolated bZip domain was used (residues 275-341) to minimize contribution from intermediate exchange originating from the C-terminal region of the bZip domain. Sequential ^1^H/^15^N/^13^C backbone assignment of the bZip domain (residue 275-351) was achieved using TROSY versions of conventional 3D triple resonance correlation experiments [HNCO, HNCA, HNCACB, HN(CO)CA, and HN(COCA)CB]^37^ at 700 MHz. Secondary ^13^C chemical shifts were calculated using Poulsen’s random coil database with corrections for pH, temperature, and neighbor amino acid correction.^38,39^ All spectra were processed using NMRPipe^40^ and displayed with SPARKY.^41^ The ^1^H–^15^N ZZ-exchange spectra were collected at 800 MHz using a TROSY variation of the ^15^N magnetization exchange pulse sequence provided by Bruker (tretexf3gpsi). The spectra were recorded at 293K with 128* (t_1_, ^15^N) × 1024* (t_2_, ^1^H) complex points and a 12820 and 2676 Hz spectral width for the ^15^N and ^1^H dimensions, respectively. The following mixing times were acquired in random order: 150, 300, 600, and 800 ms. Evidence of exchange was only observed for mixing time of 600 ms.

### Molecular dynamics simulations

All-atom MD simulations were performed for the isolated bZip domain (residues 276-351) and full-length ATF4 (1-351) using the Pronto and Nova clusters at the High-Performance Computing facility at Iowa State University. These simulations were conducted with GROMACS.2022.2 using the AMBER99SP-disp forcefield.^14^ The systems were solvated in a triclinic simulation box with TIP4P water molecules. Bonds to hydrogen were constrained via the LINCS algorithm.^39^ Short-range electrostatic and Lennard-Jones interactions were calculated with a plain coulomb cutoff of 1.0 nm. The Particle Mesh Ewald (PME) scheme with grid spacing of 0.16 nm was utilized for long range electrostatic interactions.^40^ Solvent and solute were separately coupled to a modified Berendsen thermostat (velocity rescale) with a reference temperature of 300 K and Parinello-Rahman barostat with a reference pressure of 1 bar. After energy minimization and a 1 ns equilibration, simulations were carried out using the leapfrog integration method with 2 fs timesteps. Analysis of trajectories was done using mdtraj library and in-house developed python scripts. The frequency contact maps were calculated using a 7-angstrom cut-off for backbone Ca-Ca distances.

## Supporting Information

This article contains supporting information

## Funding and Additional Information

This project is supported by funds from the Roy J. Carver Charitable Trust of Muscatine, Iowa, and from NIGMS R01 GM132561 (J.R). The content is solely the responsibility of the authors and does not necessarily represent the official views of the National Institutes of Health.

## Conflict of Interest

The authors declare that they have no conflicts of interest with the contents of this article.

## Notes

### Competing Interest Statement

The authors have declared no competing interest.

## REFERENCES

1. Ameri K., Harris A.L. (2008) Activating transcription factor 4. Int. J. Biochem Cell Biol. 40: 14–21

2. Ebert S.M. et al. (2022) Biology of activating transcription factor 4 (ATF4) and its role in the skeletal muscle atrophy. J. Nutri. 152: 926–938

3. Liang G., Hai T. (1997). Characterization of human activating transcription factor 4, a transcriptional activator that interacts with multiple domains of cAMP-responsive element-binding protein (CREB)-binding protein (CBP). J. Biol. Chem. 272: 24088–24095

4. Dey S. et al. (2012) Transcriptional repression of ATF4 gene by CCAAT/Enhancer-binding protein β (C/EBP β) differentially regulates Integrated Stress Response. J. Biol. Chem. 287: 21936–21949

5. Pakos-Zebrucka K. et al. (2016) The integrated stress response. EMBO Rep. 17: 13741395

6. Harding H.P. et al. (2000) Regulated translational initiation controls stress-induced gene expression in mammalian cells. Molecular Cell. 6: 1099–1108

7. Ebert S.M. et al. (2020) Activating transcription factor 4 (ATF4) promotes skeletal muscle atrophy by forming a heterodimer with the transcription regulator C/EBP β. J. Biol. Chem. 295: 2787–2803

8. Gill M.L. (2016) Dynamics of GCN4 facilitate DNA interaction: a model-free analysis of an intrinsically disordered region. Phys. Chem. Chem. Phys. 18: 5839–5849

9. Robustelli P. et al. (2013) Conformational Dynamics of the Partially Disordered Yeast Transcription Factor GCN4. Chem. Theory Comput. 9: 11, 5190–5200

10. Lassot I. et al. (2001) ATF4 degradation relies on a phosphorylation-dependent interaction with the SCF(betaTrCP) ubiquitin ligase. Mol Cell Biol. 21: 2192–2202

11. Ampofo E. et al. (2013) Functional interaction of protein kinase CK2 and activating transcription factor 4 (ATF4), a key player in the cellular stress response. Biochim Biophys Acta. 1833: 439–451

12. Siang S. et al. (2022) Intricate coupling between the transactivation and basic-leucine zipper domains governs phosphorylation of transcription factor ATF4 by casein kinase 2. J Biol Chem. 298:101633.

13. Du J. (2021) ATF4 promotes lung cancer cell proliferation and invasion partially through regulating Wnt/β-catenin signaling. Int J Med Sci. 18: 1442–1448

14. Ebert S.M. et al. (2020) Activating transcription factor 4 (ATF4) promotes skeletal muscle atrophy by forming a heterodimer with the transcriptional regulator C/EBPβ. J Biol Chem. 295: 2787–2803

15. Hao L. et al. (2021) ATF4 activation promotes hepatic mitochondrial dysfunction by repressing NRF1-TFAM signalling in alcoholic steatohepatitis. Gut. 70: 1933–1945

16. Freundt J. K. et al. (2018) The Transcription Factor ATF4 Promotes Expression of Cell Stress Genes and Cardiomyocyte Death in a Cellular Model of Atrial Fibrillation. Biomed Res Int. 2018: 3694362

17. Podust L.M. et al. (2001) Crystal structure of the CCAAT box/enhancer-binding protein beta activating transcription factor-4 basic leucine zipper heterodimer in the absence of DNA. J. Biol. Chem. 276 : 505–513

18. Robustelli P. et al. (2018) Developing a molecular dynamics force field for both folded and disordered protein states. Proc. Natl. Acad. Sci. USA. 115 : E4758–4766

19. Xu J. et al. (2021) Improved protein structure prediction by deep learning irrespective of co-evolution information. Nat. Mach. Intell. 3 : 601–609

20. Baumgart S. et al. (2013) Oncogenic Transcription Factors: Cornerstones of Inflammation-Linked Pancreatic Carcinogenesis. Gut. 62: 310–316

21. Good-Jacobson K.L. & Groom J.R. (2018) Tailoring Immune Responses toward Autoimmunity: Transcriptional Regulators That Drive the Creation and Collusion of Autoreactive Lymphocytes. Front. Immunol. 9: 482

22. Tutukova S. et al. (2021) The Role of Neurod Genes in Brain Development, Function, and Disease. Front. Mol Neurosci. 14: 662774

23. Mitchell S.M. & Frayling T.M. (2002) The role of transcription factors in maturity-onset diabetes of the young. Mol Genet Metab. 77: 35–43

24. Budhram-Mahadeo V.S. et al. (2021) Linking metabolic dysfunction with cardiovascular diseases: Brn-3b/POU4F2 transcription factor in cardiometabolic tissues in health and disease. Cell Death Dis 12: 267

25. Su B.G. & Henley M.J. (2021) Drugging Fuzzy Complexes in Transcription. Front Mol Biosci. 8: 795743

26. Radaeva M. et al. (2021) Drugging the ‘undruggable’. Therapeutic targeting of protein-DNA interactions with the use of computer-aided drug discovery methods. Drug Discov Today. 26: 2660–2679

27. Lao B.B. et al. (2014). In Vivo modulation of Hypoxia-Inducible Signaling by Topographical helix Mimetics. Proc. Natl. Acad. Sci. 111: 7531–7536

28. Gronemeyer, H. et al. (2004) Principles for modulation of the nuclear receptor superfamily. Nat. Rev. Drug Discov. 3: 950–964 (2004)

29. Wang P. et al. (2011) Specific blocking of CREB/DNA binding by cyclometalated platinum(II) complexes. Angew Chem Int Ed Engl. 50: 2554–2558

30. Krois A.S. et al. (2018) Long-range regulation of p53 DNA binding by its intrinsically disordered Nterminal transactivation domain. Proc Natl Acad Sci USA. 115: E11302–E11310

31. Kim J. et al. (2021) FOXO4 transactivation domain interaction with Forkhead DNA binding domain and effect on selective DNA recognition for transcription initiation. J Mol Biol. 433: 166808

32. Sun X. et al. (2021) A phosphorylation-dependent switch in the disordered p53 transactivation domain regulates DNA binding. Proc. Natl. Acad. Sci. USA 118 : e2021456118

33. Shnitkind S. et al. (2018) Structural basis for graded inhibition of CREB:DNA interactions by multisite phosphorylation Biochemistry, 57, 6964–6972

34. Perez-Borrajero C. et al. (2019) The Biophysical Basis for Phosphorylation-Enhanced DNA-Binding Autoinhibition of the ETS1 Transcription Factor. J Mol Biol. 431:593–614

35. Holmberg C.I et al. (2002) Multisite phosphorylation provides sophisticated regulation of transcription factors. Trends Biochem Sci. 27: 619–627

36. Chen M. et al. (2021) Emerging roles of activating transcription factor (ATF) family members in tumourigenesis and immunity: Implications in cancer immunotherapy. Gene Dis. 9: 981–999

37. Clore, G. M., and Gronenborn, A. M. (1998). Determining the Structures of Large Proteins and Protein Complexes by NMR. Trends Biotechnol. 16, 22–34

38. Kjaergaard, M., Poulsen, F.M. (2011) Sequence correction of random coil chemical shifts: correlation between neighbor correction factors and changes in the Ramachandran distribution J. Biomol. NMR 50, 157–165

39. Kjaergaard, M., Brander, S., Poulsen, F.M. (2011) Random coil chemical shifts for intrinsically disordered proteins: Effects of temperature and pH J. Biomol. NMR 49, 139–49

40. Delaglio, F., Grzesiel, S., Vuister, G.W., Zhu, G., Pfeifer, J., Bax, A. (1995) NMRPipe: a multidimensional spectral processing system based on UNIX pipes. J. Biomol. NMR 5, 277–293.

41. Goddard, T.D., Kneller, D.G. (2010) Sparky 3, University of California San Francisco, San Francisco, CA.

42. Hess et al. (1998) LINCS: A linear constraint solver for molecular simulations. J. Comp. Chem. 18: 1463–1472

43. Essmann et al. (1995) A smooth particle mesh Ewald method. J Chem. Phys. 103: 8577–8593.

