## Supplementary material for "Conformational landscape of the transcription factor ATF4 is dominated by disordered-mediated inter-domain coupling": Fig. S1, Fig. S2, Fig. S3

### SUPPLEMENTARY FIGURES

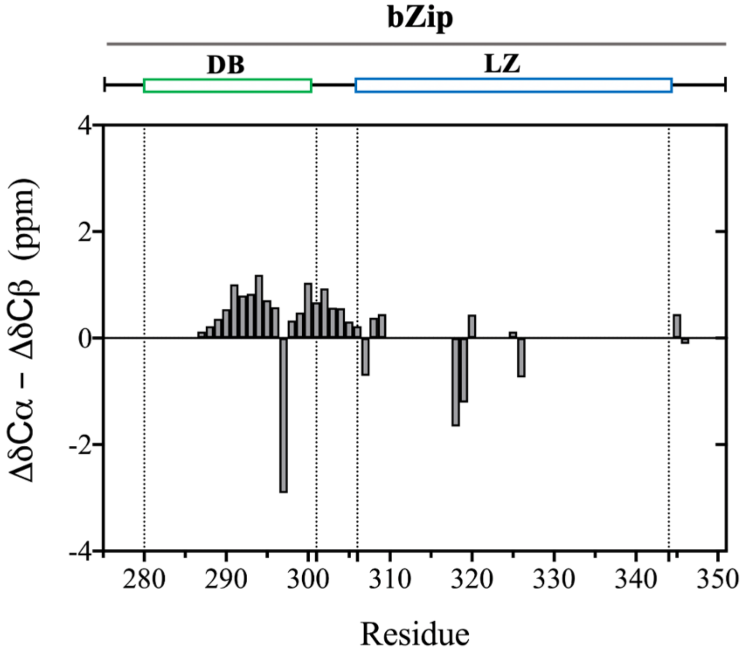


**Figure S1:** Difference in secondary Cα and Cβ chemical shifts as a function of bZip sequence. The secondary chemical shifts are calculated as the difference between the Cα and Cβ chemical shifts and chemical shifts obtained for the same residue from a random coil database. Very small deviations from zero (within 2 ppm) indicate that the isolated bZip domain is predominantly disordered in solution.

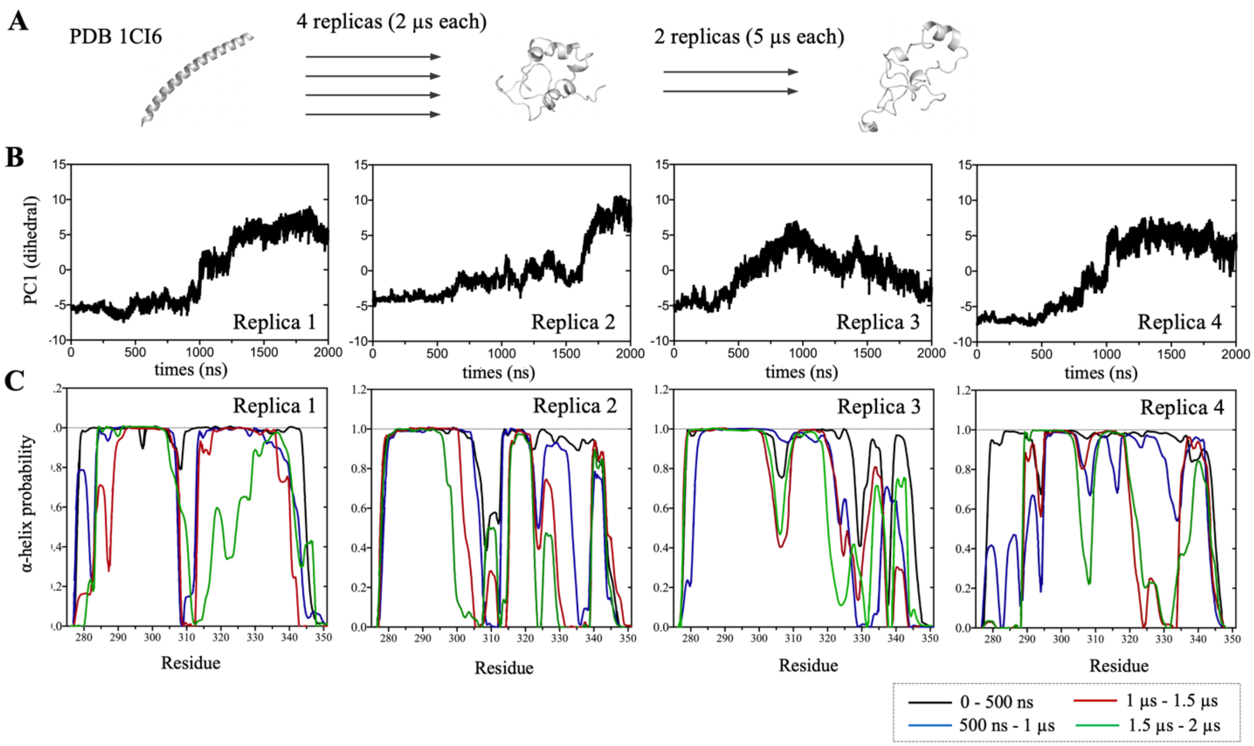


**Figure S2: (A)** Illustration of the protocol used for simulating the isolated bZip domain. Four replicas were initially set up with the original conformation provided by the reference X-ray structure (1CI6) and ran for 2 μs each. The trajectories were collectively analyzed and a new set of simulations were set up starting from a representative conformation extracted from the final run of the initial 4 replicas. Two independent replicas ran for 5 μs each were used as final production run to analyze the conformational properties of the isolated bZip domain. (**B**) Time evolution of the first principal component (PC1) in the ϕ, φ dihedral space showing equilibration of all 4 replicas within 1.5 μs. (**C**) Analysis of residue-specific α-helix probability determined for each replica over 500 ns time windows.

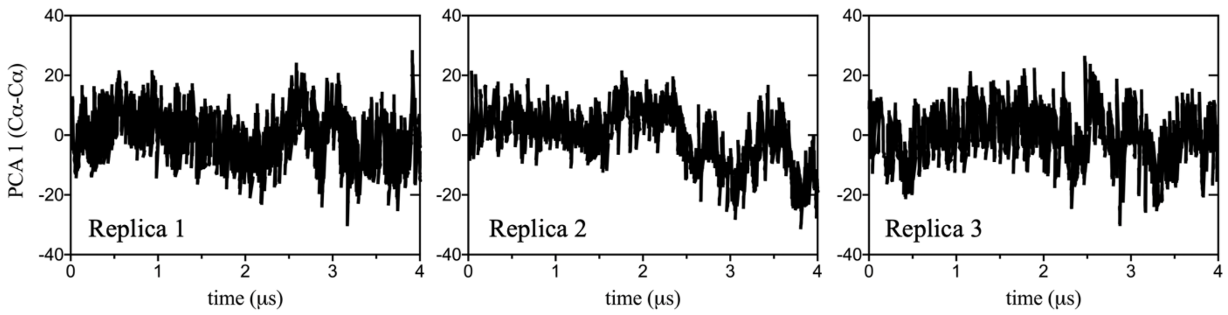


**Figure S3:** Time evolution of the first principal component (PC1) in the Cα- Cα contact space calculated for 3 independent replicas of the full-length ATF4. Evolution of PC1 showing rapid equilibration around the initial conformation.
